# Antagonistic contributions of A-type and B-type lamins to LBR localization and dynamics

**DOI:** 10.64898/2026.02.21.707184

**Authors:** Jacob Odell, Kristen Nedza, Alexander Sopilniak-Mints, Jan Lammerding

## Abstract

Lamin B receptor (LBR) is an inner nuclear membrane (INM) protein that plays crucial roles in maintaining nuclear architecture and organization of peripheral heterochromatin. Lamins and LBR both contribute to chromatin tethering at the nuclear periphery, and the expression of LBR and A-type lamins is tightly regulated during development to ensure a faithful transition between different chromatin tethering modalities. Despite its well-established association with B-type lamins, the contributions of individual lamin isoforms to LBR localization and anchorage have not been systematically examined. Here, we used mouse embryonic fibroblasts (MEFs) lacking all endogenous lamins (triple lamin knockout: TKO) to assess how specific lamin isoforms and domains regulate LBR subcellular localization and mobility. Whereas ectopic expression of either lamin B1 or lamin B2 was sufficient to tether LBR to the nuclear envelope in TKO cells, expression of lamin A increased the lateral mobility of LBR at the nuclear membrane, resulting in its displacement from the nuclear envelope to the ER. The lamin A-induced displacement of LBR was mediated by phosphorylation of LBR. Overexpression of lamin A in wild-type MEFs similarly increased LBR phosphorylation and promoted its displacement from the nuclear envelope. Collectively, these findings define isoform-specific and antagonistic roles for A-type and B-type lamins in regulating LBR anchorage at the nuclear envelope. In addition, they indicate a lamin A-dependent mechanism that may reflect a broader developmental process, since LBR and lamin A sequentially tether peripheral heterochromatin during development.

## Introduction

LBR is an integral inner nuclear membrane protein and a component of the nuclear lamina, alongside the nuclear intermediate filament lamin proteins (Worman *et al*., 1988). The name “LBR” is derived from its initial identification in a screen for lamin binding proteins, where LBR was found to have significantly higher affinity for lamin B1 (LaB1) compared to lamin A (LaA), and it was suggested that LBR may act as a “receptor” to anchor LaB1 at the inner nuclear membrane (Worman *et al*., 1988). Subsequent work showed that LaB1 can target to the nuclear membrane in the absence of LBR (Young *et al*., 2021) and no analogous “receptor” protein has been described for the A-type lamins, yet the name LBR has persisted. The N-terminal nucleoplasmic domain of LBR contains a Tudor DNA binding domain and the region responsible for binding LaB1 (Ye and Worman, 1994), whereas the C-terminus consists of several transmembrane domains and has sterol reductase activity (Holmer *et al*., 1998; Silve *et al*., 1998; Olins *et al*., 2010). According to the diffusion-retention model of inner nuclear membrane (INM) protein anchorage, INM proteins initially synthesized in the ER diffuse from the outer nuclear membrane (ONM) to the INM at nuclear pores and are subsequently retained at the INM via interactions with chromatin or nuclear envelope resident proteins (Ellenberg *et al*., 1997). This model has been extensively documented for other INM proteins such as emerin, which depends on A-type lamins for its proper subcellular localization and anchorage (Sullivan *et al*., 1999; Östlund *et al*., 1999, 2006; Odell *et al*., 2026). The diffusion-retention model has also been suggested to explain the strong anchoring of LBR to the inner nuclear membrane, perhaps through direct interactions with lamins or chromatin involving the N-terminal domain of LBR (Ellenberg *et al*., 1997).

Nonetheless, whether loss of B-type lamins perturbs LBR localization or lateral diffusion has not been directly tested previously, nor whether LaB1 and LaB2 have equal abilities to anchor LBR to the nuclear envelope. A-type lamins have also been implicated in interactions with LBR (Olins *et al*., 2010), as LaA has been identified in a detergent solubilized LBR complex isolated by immunoprecipitation (Simos and Georgatos, 1992), and a point mutation in *LMNA* (R377H) results in LBR mislocalization from the nuclear envelope (Reichart *et al*., 2004). Both LBR and lamins are known to help anchor peripheral heterochromatin in cells (Solovei *et al*., 2013; Marin *et al*., 2025), and this process is tightly regulated in development as LBR and A-type lamins sequentially tether chromatin to the nuclear periphery (Solovei *et al*., 2013), but details regarding specific contributions of individual lamins to LBR dynamics have yet to be elucidated.

Here, we systematically characterize the contributions of LaA, LaB1, and LaB2 to LBR subcellular localization and anchorage using TKO MEFs engineered to inducibly express specific lamin isoforms or lamin domains. Our studies reveal antagonistic roles of LaA and B-type lamins on LBR anchorage to the INM, with LaB1 or LaB2 being sufficient to anchor LBR at the INM and limit its lateral diffusion, whereas LaA displaces LBR from the nuclear envelope and increases LBR’s lateral diffusion when expressed in TKO MEFs. LaA induced LBR displacement was dependent on assembly of LaA into a filamentous network and mediated by phosphorylation of LBR, which promotes the translocation of LBR from the INM to the ER. Finally, we demonstrate that overexpression of LaA in wild-type MEFs is sufficient to induce LBR displacement from the INM to the ER, which we propose may reflect the process that normally occurs in development as LBR and LaA sequentially tether peripheral heterochromatin in developing tissues.

## Results

### Lamin A displaces LBR from the nuclear envelope following expression in TKO MEFs

To systematically evaluate the impact of individual lamin proteins on interactions with LBR, we took advantage of MEFs in which all three lamin genes are deleted, i.e., “triple-lamin knockout” (TKO: *Lmna*^–/–^, *Lmnb1*^–/–^, *Lmnb2*^–/–^) MEFs, and corresponding wild-type controls (Figure 1A). TKO MEFs were stably modified with expression constructs to allow for doxycycline (dox)-inducible expression of individual lamins isoforms (LaA, LaB1, LaB2) at similar expression levels (Figure 1C-D). Immunofluorescence labelling for endogenous LBR revealed that despite lacking all endogenous lamins, TKO MEFs retained LBR within the nucleus (Figure 1B), consistent with previous reports, which suggested that LBR can be targeted to the nuclear envelope and retained via interactions with chromatin, independently of lamins (Smith and Blobel, 1994; Graumann *et al*., 2007). Neither LaB1 nor LaB2 expression perturbed the nuclear localization of LBR in TKO MEFs (Figure 1E-F). Surprisingly, expression of LaA, in contrast, caused LBR to become displaced from the nuclear envelope and instead accumulate in the ER (Figure 1E), leading to a decreased ratio of nuclear-to-cytoplasmic LBR following expression of LaA (Figure 1F). These results suggest that although lamins are not required for LBR’s nuclear localization, expression of LaA, but not B-type lamins, is sufficient to displace LBR from the nuclear envelope into the ER in TKO MEFs. In wild-type cells, however, endogenous LaA expression does not cause LBR mislocalization, suggesting that the effect of LaA is dependent on the presence of B-type lamins or the ratio of LaA to B-type lamins.

**Figure 1:**
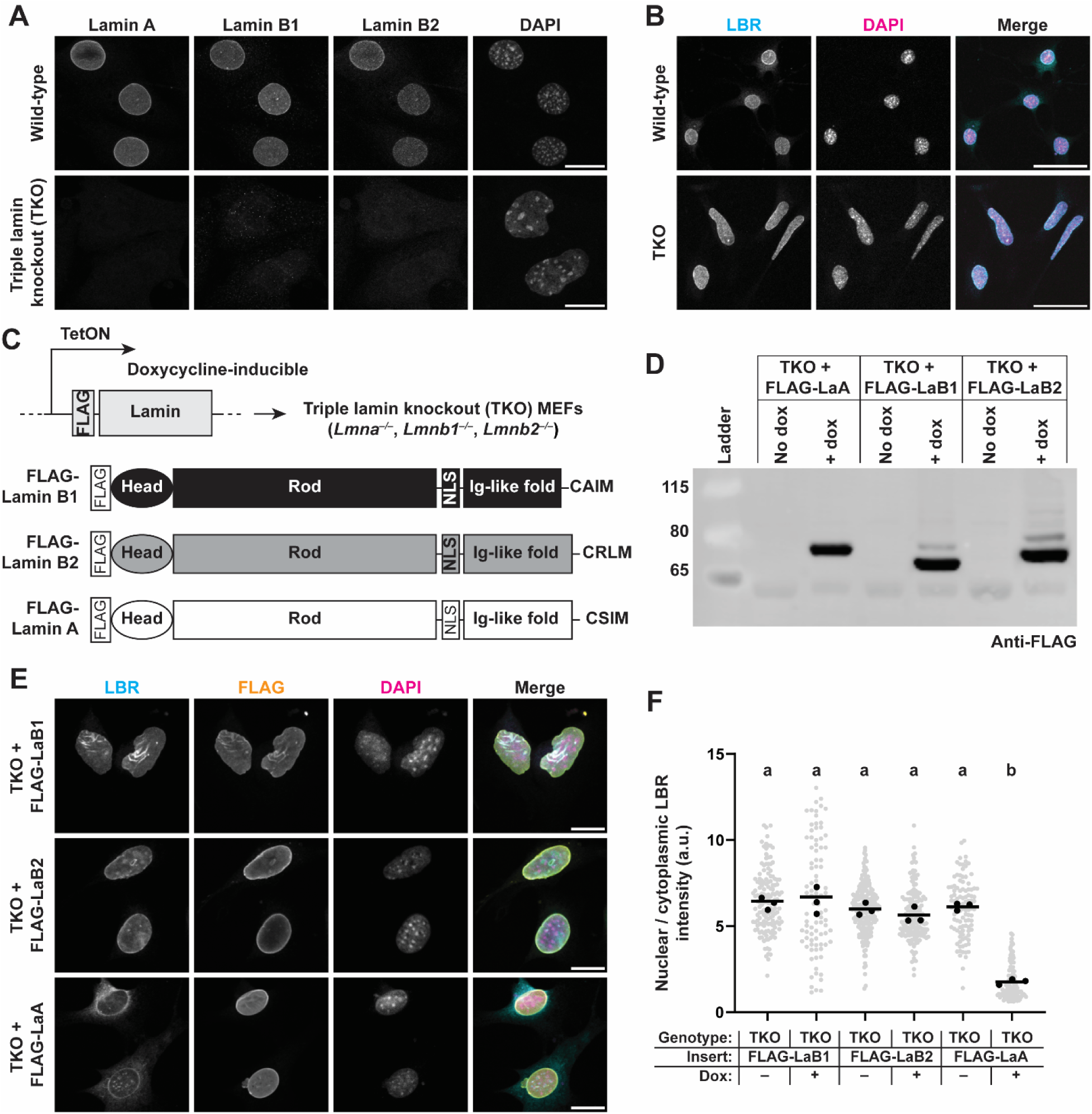
Subcellular localization of LBR in cells expressing different combinations of full-length lamins. (**A**) Validation of lamin deletion in TKO MEFs via immunofluorescence labelling for endogenous lamins. Scale bar = 20 µm. (**B**) Immunofluorescence labelling for endogenous LBR reveals nuclear enrichment in both wild-type (WT) and TKO cells. Scale bar = 50 µm. (**C**) Schematic depiction of expression strategy to re-introduce specific lamin isoforms into TKO cells. The TetON promoter allows for doxycycline-inducible expression of the desired lamin isoform. All exogenous lamin constructs include an N-terminal FLAG tag to facilitate detection with anti-FLAG antibodies. (**D**) Representative immunoblot of protein lysates from TKO cells stably modified with the indicated lamin expression construct. Equal loading was confirmed based on total protein staining. No dox conditions indicate that there is no expression of the exogenous lamin in the absence of doxycycline. (**E**) Representative immunofluorescence images of TKO cells expressing either FLAG tagged LaB1, LaB2, or LaA. Note the displacement of LBR from the nucleus following expression of LaA. Scale bar = 20 µm. (**F**) Quantification of nuclear / cytoplasmic LBR mean fluorescence intensity of cells immunofluorescently labelled as in (**E**). Where indicated, doxycycline was added at 100 ng/ml for 24 h prior to fixation and immunofluorescence labelling. All images shown are maximum intensity projections of confocal z-stacks. Grey points indicate measurements from individual cells, black points indicate replicate means, and bars indicate overall means. Different letters above each set of points represents *p* < 0.05, whereas the same letter indicates that the difference is not statistically significant, based on One-way ANOVA with Tukey’s multiple comparison test performed on the replicate means.

### B-type lamins are sufficient to anchor LBR at the INM

Given the contrasting effects of A-type and B-type lamins on LBR subcellular localization, we hypothesized that individual lamin isoforms might have different abilities to interact with LBR and to anchor it at the INM, thus influencing the lateral mobility of LBR at the nuclear envelope. A similar paradigm has been previously reported for emerin, which requires A-type lamins for effective anchoring at the INM (Östlund *et al*., 2006; Odell *et al*., 2026). To quantify the anchorage of LBR at the INM, we performed fluorescence recovery after photobleaching (FRAP) experiments using LBR-GFP transiently expressed in wild-type or TKO MEFs (Figure 2A). Compared to wild-type MEFs, TKO MEFs exhibited faster and more complete recovery of LBR-GFP following local photobleaching (Figure 2A-B), suggesting that loss of all lamins increases the lateral diffusion of LBR at the nuclear envelope, indicative of a lack of anchoring at the nuclear membrane. To determine the ability of each individual lamin isoform to anchor LBR and restrict the lateral diffusion of LBR at the nuclear envelope, we performed corresponding FRAP experiments with TKO MEFs expressing either LaA, LaB1, or LaB2. Expression of LaB1 or LaB2 was sufficient to completely restore wild-type levels of LBR mobility in TKO MEFs (Figure 2B-C), indicating that either B-type lamin can effectively anchor LBR. In contrast, LaA expression *increased* the mobility of LBR-GFP in TKO MEFs (Figure 2D), consistent with the loss of LBR from the nucleus in our immunofluorescence labelling for endogenous LBR in the TKO MEFs expressing LaA (Figure 1E, F). Because the mobility of LBR-GFP was extremely low, we were unable to robustly calculate median recovery half times (*t*_1/2_) and instead present the LBR-GFP mobile fraction for quantitative comparisons of the FRAP data. Quantification of the mobile fraction of LBR-GFP confirmed that expression of LaB1 or LaB2 in TKO MEFs restored LBR mobility to wild-type levels, whereas expression of LaA in TKO MEFs increased the mobile fraction of LBR-GFP (Figure 2E). Collectively, these results suggest opposing effects of LaA and B-type lamins on LBR localization and anchorage, with B-type lamins acting to help tether LBR to the INM and reduce its lateral diffusion, and LaA inducing LBR displacement and increasing LBR mobility within the nuclear membranes in TKO MEFs.

**Figure 2:**
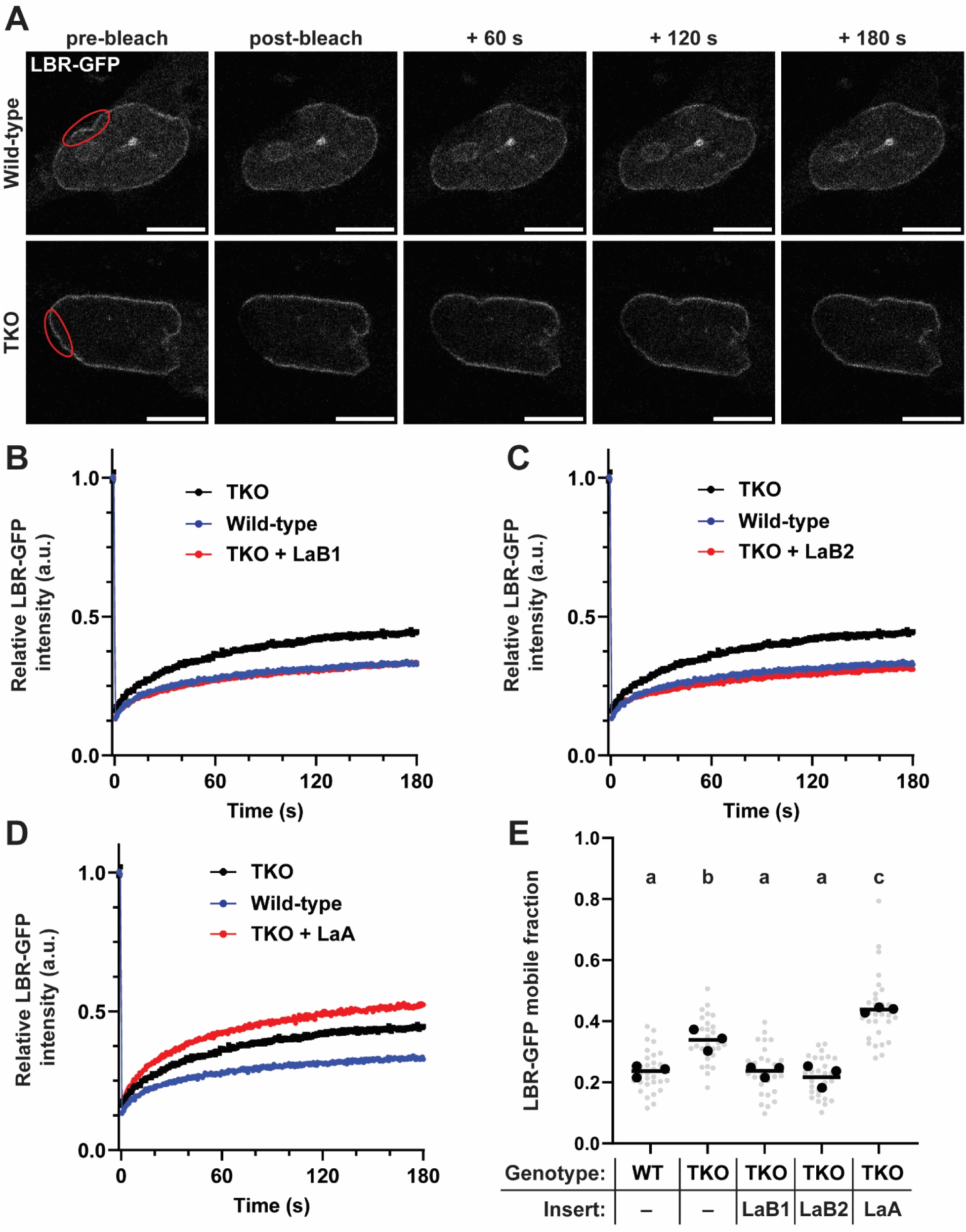
Mobility of LBR-GFP in cells expressing different combinations of full-length lamins. (**A**) Representative image series of FRAP experiment on LBR-GFP expressed in wild-type or TKO MEFs. Red outline indicates the target bleach region. Scale bar = 10 µm. (**B-D**) Quantification of LBR-GFP mean fluorescence intensity within the bleached region over time for TKO MEFs expressing either LaB1 (**B**), LaB2 (**C**), or LaA (**D**). Data are normalized such that the average prebleach intensity for each cell equals 1. Error bars indicate mean ± s.e.m. (**E**) Quantification of the LBR-GFP mobile fraction, based on the proportion of LBR-GFP intensity that recovers after 180 s relative to the prebleach and postbleach intensities for each cell. Grey points indicate measurements from individual cells, black points indicate replicate means, and bars indicate overall means. Different letters above each set of points represents *p* < 0.05, whereas the same letter indicates that the difference is not statistically significant, based on One-way ANOVA with Tukey’s multiple comparison test performed on the replicate means.

We next considered whether LaA-induced LBR displacement to the ER requires nuclear envelope breakdown in mitosis. One possible explanation for the LaA-induced LBR displacement is that LBR is unable to integrate into the assembling nuclear lamina following the completion of mitosis. Alternatively, LBR may be displaced during interphase, concurrently with LaA expression and assembly into the lamina, without nuclear envelope breakdown. To distinguish between these possibilities, we performed live cell imaging of TKO MEFs constitutively expressing LBR-GFP and with inducible expression of LaA. Monitoring intracellular LBR-GFP distribution following induction of LaA expression revealed that LBR displacement occurred in interphase, i.e., even in the absence of cell division (Supplemental Figure 1). These results suggest that nuclear envelope breakdown and reformation is not a prerequisite for LaA-induced LBR displacement.

### Full-length B-type lamin is required to effectively anchor LBR at the INM

To better understand how LaA and B-type lamins differentially modulate LBR localization and anchorage, we designed lamin truncations to explore which specific domains of B-type lamins were required to anchor LBR at the nuclear envelope. All lamins have a highly conserved tripartite domain structure consisting of a small N-terminal head, central coiled-coil rod domain, and a C-terminal tail that contains a nuclear localization signal (NLS) and an immunoglobulin-like (Ig-like) fold (Aebi *et al*., 1986; Stick and Peter, 2020). Since LaB1 and LaB2 were equally able to anchor LBR at the INM and thus restrict its mobility (Figure 2B-C), we limited experiments to LaB1 as a representative B-type lamins and used truncations of LaB1 for subsequent experiments. Truncated lamins were designed to allow for expression of either the LaB1 Head + Rod domains (LaB1^H+R^) or the LaB1 Tail domain (LaB1^Tail^) in TKO MEFs (Figure 3A). Both expression constructs included the LaB1 NLS to allow for import of the truncated lamins into the nucleus. Accordingly, both LaB1^H+R^ and LaB1^Tail^ localized to the nucleus based on immunofluorescence staining for the FLAG tag attached to the N-terminus of each construct (Figure 3B). Neither of the LaB1^H+R^ or LaB1^Tail^ LaB1 truncations altered the subcellular localization of endogenous LBR (Figure 3B-C), consistent with the results from full-length LaB1 (Figure 1E-F). We performed FRAP on LBR-GFP in TKO MEFs expressing truncated LaB1 constructs to determine whether either truncation affected the mobility of LBR at the nuclear envelope. Although the recovery curves of LBR-GFP in TKO cells expressing either the LaB1^H+R^ or LaB1^Tail^ truncations fall between those of the wild-type and TKO controls (Figure 3D-E), quantification of the LBR-GFP mobile fraction did not reveal a statistically significant difference in LBR-GFP mobility between TKO controls and TKO MEFs expressing either LaB1^H+R^ or LaB1^Tail^ (Figure 3F), in contrast to full-length LaB1, which restored wild-type levels of LBR-GFP mobility (Figure 2B). These results indicate that although the LaB1 truncations may have subtle effects on the dynamics of LBR-GFP mobility, neither LaB1 truncation was sufficient to restore wild-type levels of diffusion, which contrasts with the effect of the full-length LaB1 construct (Figure 1E-F). These results suggest that full-length LaB1 is required for effective anchorage of LBR to the nuclear envelope in MEFs.

**Figure 3:**
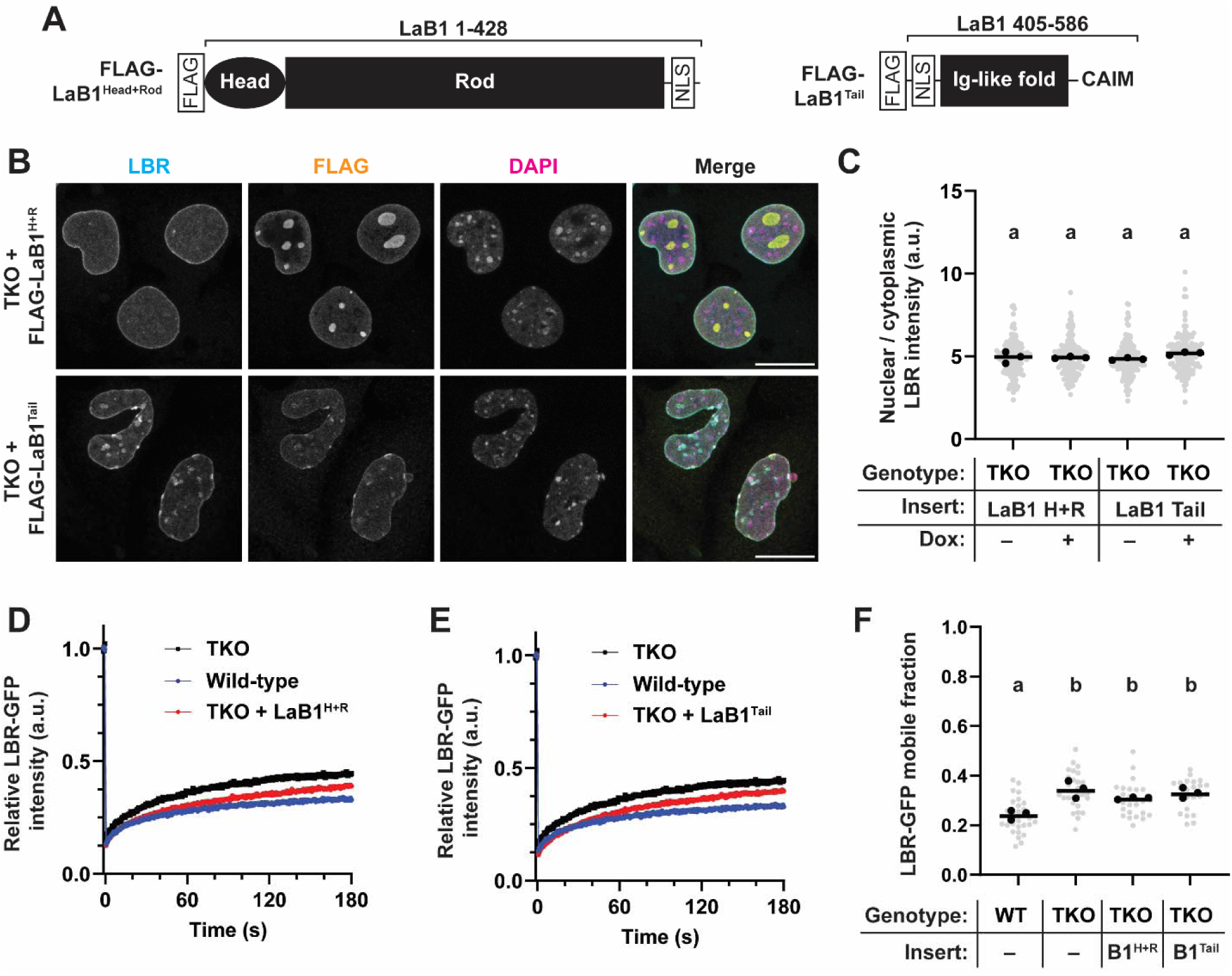
Lamin B1 (LaB1) truncations are unable to tether LBR to the INM in TKO MEFs. (**A**) Schematic depiction of the LaB1 truncations. The amino acids derived from LaB1 are indicated above each construct. (**B**) Immunofluorescence labelling for LBR and FLAG-tagged lamin truncations expressed in TKO MEFs. Scale bar = 20 µm. (**C**) Quantification of nuclear/cytoplasmic LBR immunofluorescence intensity, as in Figure 3B. (**D-E**) LBR-GFP recovery curves after photobleaching in TKO MEFs expressing either LaB1^H+R^ (**D**) or LaB1^Tail^ truncation (**E**). (**F**) LBR-GFP mobile fraction quantification in TKO cells expressing LaB1 truncations. For (**C**) and (**F**), grey points indicate measurements from individual cells, black points indicate replicate means, and bars indicate overall means. Different letters above each set of points represents *p* < 0.05, whereas the same letter indicates that the difference is not statistically significant, based on One-way ANOVA with Tukey’s multiple comparison test performed on the replicate means.

### Full-length LaA and incorporation of LaA into the lamina are requirements to displace LBR from the nuclear envelope in TKO MEFs

To explore which domains of LaA are responsible for the displacement of LBR from the nuclear envelope to the ER in TKO cells, we generated truncations of LaA containing the head + rod domains (LaA^H+R^) or the tail domain (LaA^Tail^), analogous to those described for LaB1 (Figure 4A). As with the LaB1 truncations, both LaA truncation constructs localized to the nucleus, but neither the LaA^H+R^ nor the LaA^Tail^ truncation were able to displace LBR from the INM (Figure 4B-C), unlike full-length LaA (Figure 1E-F). This suggests that full-length LaA is necessary to displace LBR from the nuclear envelope. To probe whether assembly of LaA into filaments at the nuclear lamina is required for LBR displacement, we expressed designed ankyrin repeat proteins (DARPins) that selectively bind to A-type lamins and prevent their incorporation into the peripheral nuclear lamina (Zwerger *et al*., 2015). The LaA specific DARPin (D-LaA_1) has been used previously to explore the requirement of lamin assembly on emerin anchorage at the nuclear envelope (Odell *et al*., 2026). Expression of D-LaA_1 prevented LBR displacement following expression of full-length LaA (Figure 4D-E). Expression of a control DARPin (D-E3.5) that does not bind to LaA and thus does not affect its assembly (Zwerger *et al*., 2015; Odell *et al*., 2026) did not prevent LaA peripheral localization nor displacement of LBR following LaA expression (Figure 4D-E). These results indicate that not only is full-length LaA required to displace LBR, but LaA must assemble into a filamentous nuclear lamina, suggesting that the assembled LaA network is required to displace LBR in TKO MEFs.

**Figure 4:**
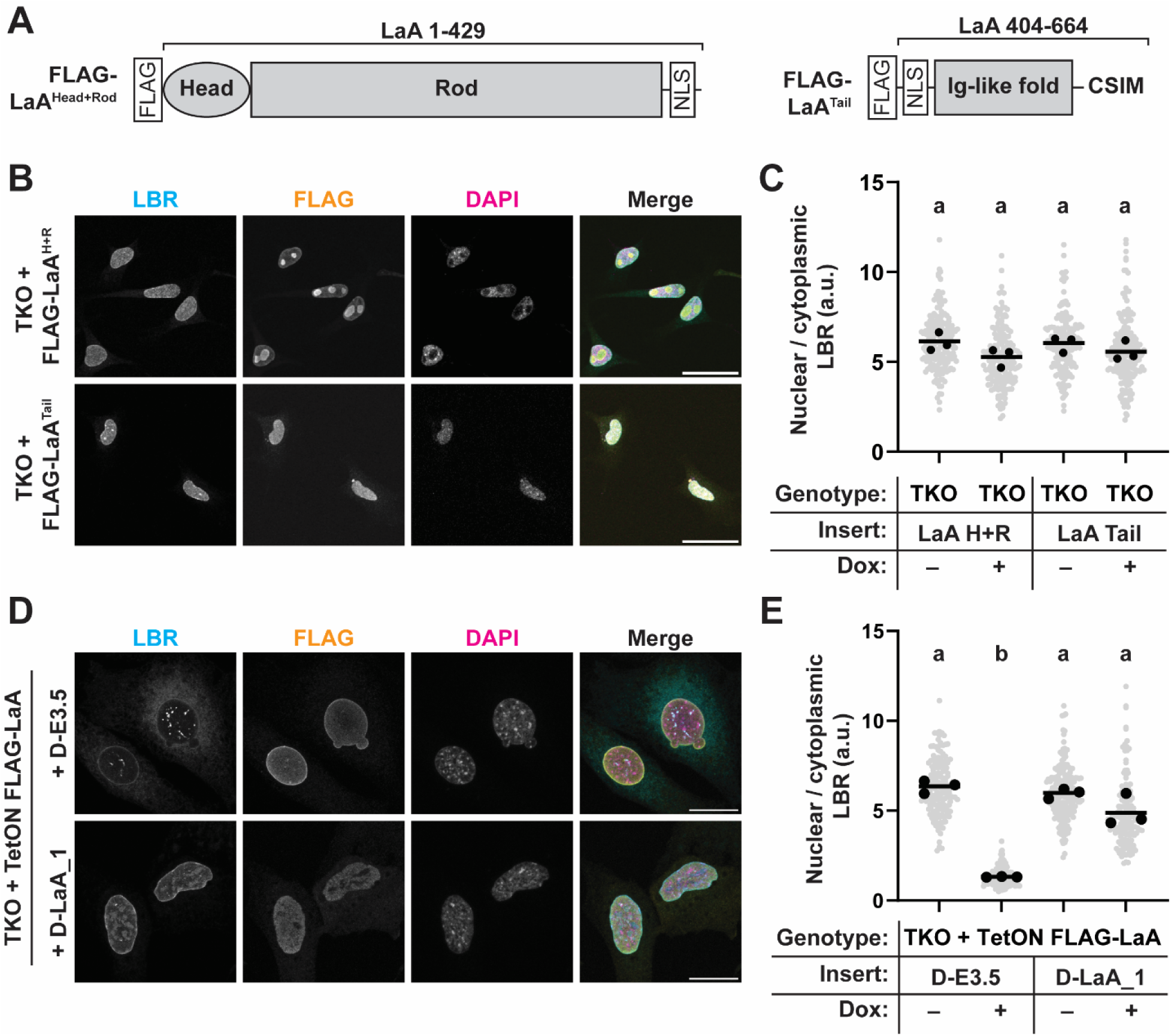
LBR is displaced by full length, assembled LaA in TKO MEFs. (**A**) Schematic depiction of the LaA^H+R^ and LaA^Tail^ truncations. The amino acids derived from LaA are indicated above each construct. (**B**) Immunofluorescence labelling for LBR and FLAG-tagged lamin truncations expressed in TKO MEFs. Scale bar = 50 µm. (**C**) Quantification of nuclear/cytoplasmic LBR immunofluorescence intensity of the cells in panel (**B**). (**D**) Immunofluorescence labelling for LBR and FLAG-LaA in TKO cells expressing DARPins. The control DARPin (D-E3.5) does not interfere with LaA assembly (Zwerger *et al*., 2015), whereas the experimental DARPin D-LaA_1 prevents the incorporation of LaA into the lamina network, evidenced by the loss of FLAG signal at the nuclear periphery. Scale bar = 10 µm. (**E**) Quantification of nuclear/cytoplasmic LBR immunofluorescence intensity of TKO MEFs expressing DARPins and FLAG-LaA. For (C) and (E), grey points indicate measurements from individual cells, black points indicate replicate means, and bars indicate overall means. Different letters above each set of points represents *p* < 0.05, whereas the same letter indicates that the difference is not statistically significant, based on One-way ANOVA with Tukey’s multiple comparison test performed on the replicate means.

### LaA expression in TKO cells induces LBR phosphorylation to drive displacement from the nuclear envelope

We next aimed to understand the mechanism by which LaA expression results in LBR displacement in TKO MEFs. LBR is phosphorylated by several families of kinases, including serine-arginine protein kinases (SRPKs) and cyclin-dependent kinases (CDKs) (Nikolakaki *et al*., 1996; Papoutsopoulou *et al*., 1999; Takano *et al*., 2002). Inhibiting these kinases pharmacologically results in an increase in nuclear LBR, whereas decreased expression of multiple phosphatases results in increased cytoplasmic localization of LBR (Mimura *et al*., 2016). Thus, phosphorylation of LBR leads to its enrichment in the cytoplasm, whereas dephosphorylation of LBR promotes its localization to the nucleus. We hypothesized that LaA expression could lead to activation or recruitment of a kinase at the nuclear envelope to phosphorylate LBR, which could explain its translocation from the nuclear envelope to the ER in the TKO MEFs. To assess the phosphorylation status of LBR, we used Phos-tag gels, which allow for the separation of a protein of interest based on its phosphorylation levels via binding of the Phos-tag reagent to negatively charged phosphates on proteins which reduces protein mobility during electrophoresis, resulting in discrete bands that correspond to differently phosphorylated species (Kinoshita *et al*., 2006). We first demonstrated that LaA expression did not significantly alter the total protein levels of LBR (Figure 5A-B), even after 72 h of LaA expression, indicating that LBR is not degraded once it is displaced from the nuclear envelope. Separation of lysates from TKO MEFs expressing LaA via Phos-tag gels revealed an increase in highly phosphorylated LBR following LaA expression compared to the no dox controls (Figure 5C-D), supporting our hypothesis that LBR displacement is driven by increased phosphorylation of LBR.

**Figure 5:**
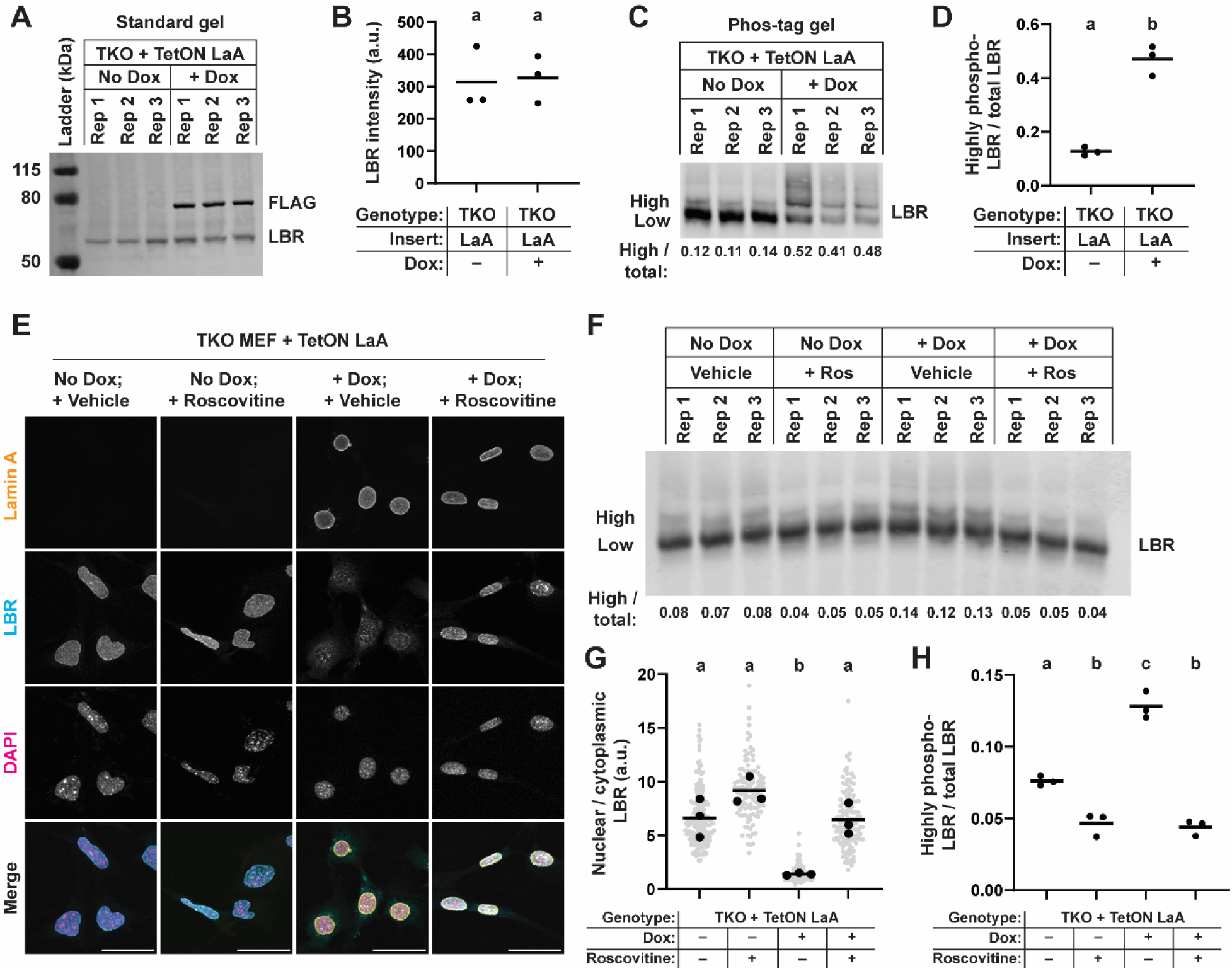
LaA expression drives CDK-mediated LBR phosphorylation to displace LBR from the nuclear envelope. (**A**) Immunoblot of lysates from TKO MEFs treated with no dox or 100 ng/ml doxycycline for 72 h prior to lysis, probed with antibodies recognizing endogenous LBR or the FLAG epitope tag on LaA. (**B**) Quantification of LBR band intensity in (**A**), normalized to total protein in each lane based on Coomassie staining of the gel after transfer. (**C**) Lysates from TKO MEFs expressing or not expressing LaA were separated by Phos-tag gel to resolve differently phosphorylated forms of LBR. Numbers below each lane quantify the proportion of highly phosphorylated LBR (upper band) divided by the total LBR signal in each lane. (**D**) Graphical representation of data in (C). (**E**) Representative images of cells treated with no dox or 100 ng/ml doxycycline for 24 h to induce LaA expression, then treated with vehicle (DMSO) or 20 µM roscovitine for an additional 24 hours. Maximum projections of confocal z-stacks are shown. Note the displacement of LBR following LaA expression and vehicle treatment, which is reversed following treatment with roscovitine. Scale bar = 50 µm. (**F**) Lysates from conditions described in (**E**) were separated by Phos-tag gel, and the proportion of highly phosphorylated LBR / total LBR was calculated as in (**C**). (**G**) Quantification of nuclear/cytoplasmic LBR intensity of cells described in (**E**). (**H**) Quantification of Phos-tag experiment described in (**F**). For (**B**) and (**D**), points represent replicate means and bars indicate overall means. Different letters over each set of points indicate *p* < 0.05, whereas the same letter indicates that the difference was not statistically significant based on two-tailed unpaired *t*-test. For (**G**) and (**H**), grey points indicate measurements from individual cells, black points indicate replicate means, and bars indicate overall means. Different letters above each set of points represents *p* < 0.05, whereas the same letter indicates that the difference is not statistically significant, based on One-way ANOVA with Tukey’s multiple comparison test performed on the replicate means.

Since LaA expression led to increased LBR phosphorylation and LBR translocation into the ER, we decided to modulate LBR phosphorylation by pharmacologically inhibiting kinases responsible for LBR phosphorylation. We first tested whether inhibiting SRPKs, which are known to phosphorylate LBR, could reverse LBR displacement, but found that treatment with the SRPK inhibitor SRPIN340 failed to restore the normal nuclear localization of LBR following expression of LaA (Supplemental Figure S2). As an alternative approach, we treated TKO MEFs expressing either LaA or no-dox controls with roscovitine, an inhibitor of CDK kinases, including CDK1, CDK2, CDK5, CDK7, and CDK9 (Bach *et al*., 2005), which has been previously shown to prevent LBR phosphorylation and increase the fraction of nuclear LBR (Mimura *et al*., 2016). Roscovitine treatment resulted in a decreased fraction of highly phosphorylated LBR in TKO cells and TKO cells expressing LaA (Figure 5F-H), indicating that roscovitine successfully blocked LBR phosphorylation. Notably, roscovitine treatment was sufficient to reverse the LBR displacement following LaA expression and led to enrichment of LBR in the nucleus in TKO cells expressing LaA (Figure 5E, G). In contrast, LBR was displaced from the nuclear envelope to the ER in vehicle-only treated TKO cells expressing LaA (Figure 5E, G), matching our earlier results in non-treated TKO cells expressing LaA (Figure 1E-F). Taken together, these results suggest that LaA displaces LBR from the nuclear envelope in a CDK-mediated LBR phosphorylation-dependent manner, since the displacement could be reversed by inhibiting CDKs.

### Overexpression of LaA or *LMNA* R377H mutation displaces LBR from the nuclear envelope in wild-type cells

Previous work has described a *LMNA* mutation (R377H) that causes Emery-Dreifuss muscular dystrophy (EDMD) and that is unique among studied *LMNA*-EDMD mutations in that patient cells harboring this mutation display LBR displaced into the cytoplasm and ER (Reichart *et al*., 2004). To validate these prior results, we designed an expression construct for LaA^R377H^ (Figure 6A) and expressed this construct in wild-type MEFs to mimic the heterozygous presentation of this mutation in patients, i.e., the mutant LaA is expressed in combination with at least one healthy, non-mutated copy of LaA. As an important control, we also overexpressed wild-type LaA (LaA^WT^) in the wild-type MEFs. Interestingly, overexpression of either LaA^WT^ or LaA^R377H^ resulted in displacement of endogenous LBR from the nuclear envelope to the ER (Figure 6B-C). Expression of either LaA^WT^ or LaA^R377H^ also resulted in an increased pool of highly phosphorylated LBR (Figure 6D-E), consistent with our proposed phosphorylation-dependent model of LBR displacement from the nuclear envelope.

**Figure 6:**
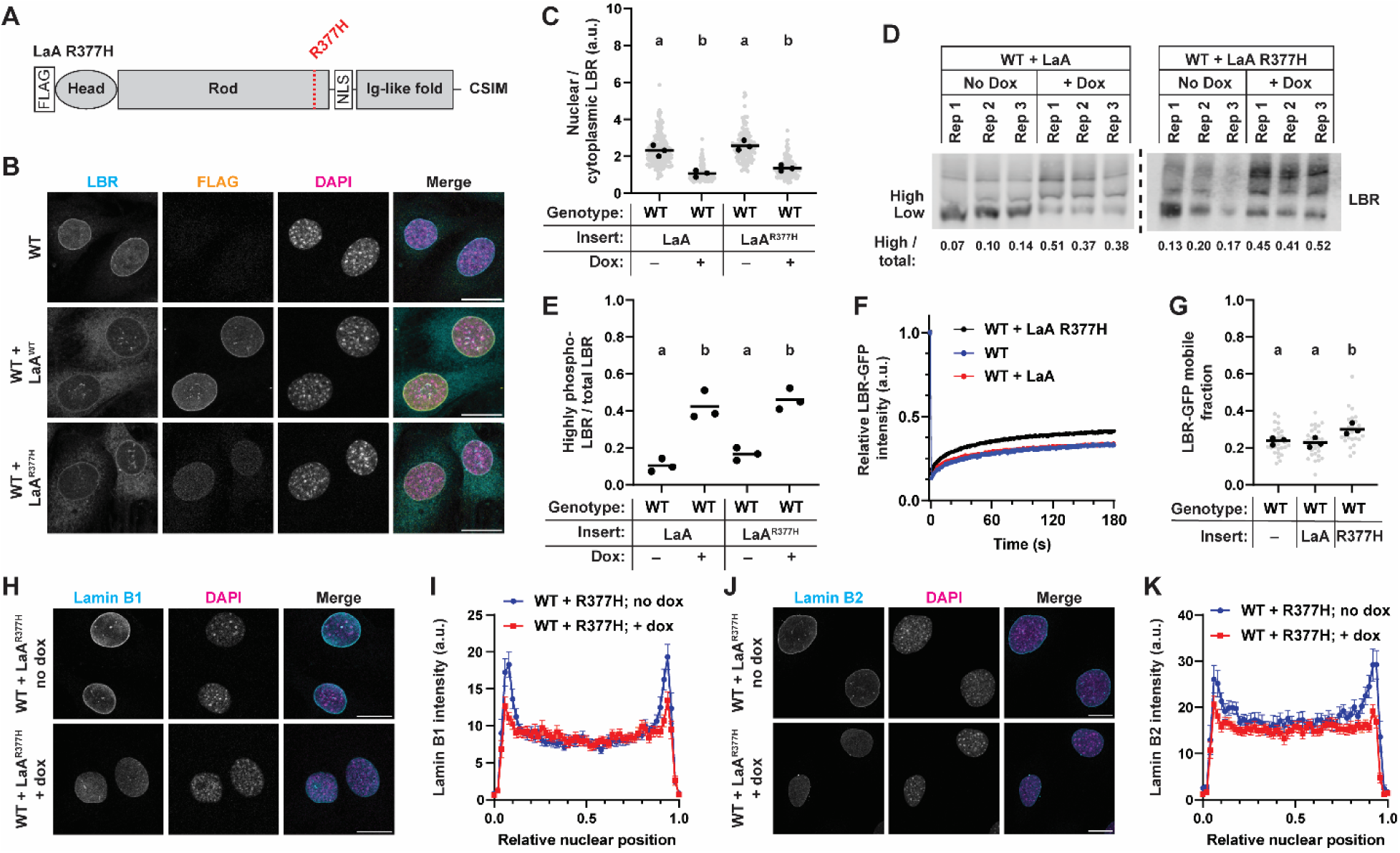
LaA^R377H^ displaces LBR from the nuclear envelope and perturbs the assembly of LaB1 into the lamina. (**A**) Schematic depiction of the location of the LaA R377H mutation, which is at the end of coil 2 of the lamin rod domain. (**B**) Immunofluorescence labeling for LBR in wild-type cells expressing the LaA^WT^ or LaA^R377H^ mutation reveals displacement of LBR from the nucleus into the cytoplasm and ER. Scale bar = 10 µm. (**C**) quantification of nuclear/cytoplasmic LBR intensity of cells described in (**B**). (**D-E**) Lysates from wild-type MEFs expressing LaA^WT^ (left) or LaA^R377H^ (right) were separated by Phos-tag gel (**D**), and the ratio of highly phosphorylated / total LBR was quantified (**E**). (**F-G**) LBR-GFP FRAP recovery curves (**F**) and mobile fraction quantification (**G**) in wild-type MEFs expressing LaA^WT^ or LaA^R377H^. (**H**) Immunofluorescence labelling for endogenous LaB1 in wild-type MEFs expressing LaA^R377H^. Scale bar = 10 µm. (**I**) Intensity profiles of LaB1 measured by drawing a line through the midplane of the nucleus. Peaks at the edges of the plot represent the nuclear envelope signal. Points and error bars indicate mean ± s.e.m. (**J**) Immunofluorescence labelling for endogenous LaB2 in wild-type MEFs expressing LaA^R377H^. Scale bar = 10 µm. (**K**) Intensity profiles of LaB2 measured by drawing a line through the midplane of the nucleus. Peaks at the edges of the plot represent the nuclear envelope signal. Points indicate mean ± s.e.m. For (**C**) and (**G**), grey points represent measurements from individual cells, black points represent replicate means and bars indicate overall means. For (**E**), points indicate measurements from individual replicates and bars indicate overall means. Different letters above each set of points represents *p* < 0.05, whereas the same letter indicates that the difference is not statistically significant, based on One-way ANOVA with Tukey’s multiple comparison test performed on the replicate means.

However, although both LaA^WT^ and LaA^R377H^ led to increased LBR phosphorylation and displacement from the nucleus in wild-type MEFs, we observed different effects of LaA^WT^ and LaA^R377H^ on LBR-GFP mobility at the nuclear envelope. Overexpression of LaA^WT^ in wild-type MEFs had no effect on LBR-GFP mobility compared to parental wild-type cells, whereas LaA^R377H^ overexpression led to increased LBR-GFP mobility at the nuclear envelope (Figure 6F-G). These results suggest that overexpression of LaA^WT^ drives LBR phosphorylation and displacement in wild-type MEFs without perturbing the mobility of LBR-GFP. Given our previous data demonstrating the importance of B-type lamins in anchoring LBR and restricting its lateral diffusion, we considered whether LaA^R377H^ may perturb the networks formed by endogenous B-type lamins. Whereas overexpression of LaA^WT^ in wild-type MEFs had no effect on the peripheral localization of LaB1 or LaB2 (Supplemental Figure 3A-D), expression of LaA^R377H^ dominantly perturbed the subnuclear localization of LaB1 (Figure 6H-I) and LaB2 (Figure 6J-K) by reducing the nuclear peripheral signal following immunofluorescence labelling.

Based on these results, we propose that the LaA^R377H^ mutation not only drives LBR phosphorylation and displacement but also increases the mobility of LBR-GFP by interfering with B-type lamin networks, which may hinder the ability of B-type lamins to tightly anchor LBR to the nuclear envelope. Taken together, these data highlight that the subcellular localization and membrane anchorage of LBR are not always correlated. Subcellular localization of LBR is primarily determined by phosphorylation, consistent with previous reports (Mimura *et al*., 2016), as overexpression of LaA results in increased LBR phosphorylation and translocation into the ER. However, even when LBR is highly phosphorylated, some LBR is still retained at the nuclear envelope, and the mobility of this nuclear envelope-associated pool of LBR is dependent on the presence and assembly of B-type lamins into the nuclear lamina.

## Discussion

Here, we present a systematic characterization of how specific lamin isoforms contribute to the subcellular localization and lateral mobility of LBR at the nuclear envelope. Our data highlight that the subcellular localization and anchorage of LBR at the nuclear envelope are not always correlated, since TKO MEFs display normal LBR localization but have increased LBR mobility compared to wild-type cells, and wild-type cells overexpressing LaA have aberrant LBR localization but normal LBR mobility at the nuclear envelope. Based on our experimental results, we postulate that the subcellular localization of LBR is most strongly determined by LBR’s phosphorylation status, consistent with previous reports (Mimura *et al*., 2016). Increased expression of LaA results in increased LBR phosphorylation that promotes LBR translocation from the nuclear envelope to the ER. The mobility of LBR at the nuclear envelope is most strongly determined by the presence of an intact network of B-type lamins, given that TKO MEFs display increased mobility of LBR-GFP compared to wild-type cells, which can be rescued by reintroduction of full-length LaB1 or LaB2. In wild-type cells with physiological expression levels of LaA and B-type lamins, LBR exhibits nuclear localization and is strongly anchored to the nuclear envelope, suggesting that a balance exists between A-type and B-type lamins that promotes LBR nuclear localization and anchorage. Perturbations to this balance can alter LBR subcellular localization and/or anchorage to the nuclear envelope.

Consistent with a diffusion retention model where B-type lamins act to anchor LBR at the nuclear envelope, we find that full-length B-type lamins are required for this function. Additionally, full-length LaA is required to displace LBR in TKO cells, suggesting that the emergent networks of LaA versus LaB1 contribute to these proteins’ abilities to modulate LBR. These data are further supported by the results with the LaA^R377H^ mutation that perturbed LaB1 and LaB2 localization at the nuclear rim, which is predicted to abrogate the ability of B-type lamins to interact with and anchor LBR at the nuclear envelope. Future work will be needed to determine whether altered LBR anchorage contributes to disease phenotypes associated with the R377H mutation, or whether LBR anchorage represents a secondary consequence of broader lamina disruption.

The displacement of LBR from the nuclear envelope following LaA overexpression may recapitulate features of a transition that occurs during mammalian development, as LBR and LaA sequentially tether peripheral heterochromatin in developing tissues (Solovei *et al*., 2013). Lamin A/C is lowly expressed early in development, whereas LBR is initially enriched in many tissues and helps to anchor peripheral heterochromatin to the nuclear envelope (Solovei *et al*., 2013). Lamin A/C expression correlates with a downregulation of LBR, as the proteins swap roles in organizing peripheral heterochromatin in several tissues (Solovei *et al*., 2013). We speculate that LaA expression promotes LBR phosphorylation, potentially via recruitment or activation of CDKs at the nuclear envelope, thereby reducing LBR-chromatin interactions and favoring engagement of LaA-chromatin tethers. Of interest, previous work demonstrated an increased affinity of LaB1 for phosphorylated LBR relative to non-phosphorylated LBR (Appelbaum *et al*., 1990). Since LaB1 is ubiquitously expressed in normally developing tissues, LaB1 would be predicted to retain phosphorylated LBR within the nucleus and prevent translocation into the cytoplasm. However, in the TKO MEFs, which lack LaB1 and LaB2, there are no B-type lamins present to anchor LBR at the nuclear envelope following LaA expression, allowing for the increased mobility and displacement of LBR into the cytoplasm and ER. Future work will be needed to test these hypotheses, perhaps by examining the localization of LaA and LBR during development in mouse models lacking LaB1 (Vergnes *et al*., 2004) or both B-type lamins (Kim *et al*., 2011).

The present study has some limitations. Although the roscovitine treatment experiments suggest that LaA-driven LBR displacement is mediated by CDK phosphorylation, it remains to be determined how exactly LaA activates or recruits kinases to phosphorylate LBR, and which of the several kinases inhibited by roscovitine is involved. This is a challenging question to answer, as candidate kinases identified via affinity or proximity-based assays may be acting on LaA itself, and LaA is phosphorylated by several CDKs that assist in lamina disassembly at the onset of mitosis. An additional limitation is that all of the exogenous expression constructs used in this study contain an N-terminal FLAG tag, which has been shown previously to slightly impair lamin function (Odell and Lammerding, 2024). By using the same small tag for all expression constructs, we hope that any biases are uniformly introduced and do not expect this to alter our conclusions.

Taken together, this work defines novel isoform-specific roles of A-type and B-type lamins in regulating LBR anchorage and its mobility at the nuclear envelope. These results have broad impact, given the important role of lamins and LBR in organizing chromatin at the nuclear periphery in different cell types, and provide a mechanistic basis for how shifts in nuclear lamina composition can reconfigure the organization of inner nuclear membrane proteins.

## Methods

### Cell culture

Wild-type and *Lmna*^–/–^ MEFs were a kind gift of Colin Stewart (Sullivan *et al*., 1999). Triple lamin knockout MEFs were a kind gift from Stephen Young and Loren Fong and have been described in depth previously (Chen *et al*., 2018, 2021). Cells were maintained in DMEM supplemented with 10% FBS and 1% penicillin/streptomycin in a humidified incubator at 37°C and 5% CO_2_. Cells were passaged at 80-90% confluency and routinely checked for mycoplasma contamination by PCR. To induce expression of exogenous lamins, 100 ng/ml doxycycline was added to media for 24 h prior to experiments. For inhibitor treatment experiments, SRPIN340 (Cell signaling #62630) was dissolved in DMSO and added to cells at a final concentration of 10 µM. Roscovitine (Sigma R7772) was dissolved in DMSO and added to cells at a final concentration of 20 µM.

### Genetic construct information

Expression constructs for full-length LaA and LaB1 and the lamin truncations have been described previously (Odell *et al*., 2025, 2026). Transient expression constructs for mCherry-DARPins were a kind gift of Ohad Medalia (Zwerger *et al*., 2015). The LaB2 construct was cloned via Gibson assembly following PCR amplification of mLmnb2-pCMV (a kind gift from Loren Fong), and digestion of the pPB-rtTA-hCas9-puro-PB backbone (Wang *et al*., 2017) using AgeI and NheI. The stable expression vector for LBR-GFP was cloned via Gibson assembly from two existing plasmids. For the backbone, pLVX-EF1a-EGFP-Emerin-IRES-Hygromycin, a kind gift from David Andrews (Addgene plasmid # 134864) (Schormann *et al*., 2020) was digested with NotI and BamHI. Full length LBR-GFP was amplified via PCR using “LBR-pEGFP”, a kind gift from Eric Schirmer (Addgene plasmid # 61996) (Zuleger *et al*., 2011). Note that full length LBR was used, rather than previously described LBR-GFP constructs that contain the N-terminus and first transmembrane domain of LBR (Ellenberg *et al*., 1997; Östlund *et al*., 2006). A table of primers used for PCR amplification of these fragments is described in Supplemental Table 1. Following Gibson assembly and plasmid purification, successful cloning was verified by Sanger sequencing of the insert. For modification with the dox-inducible LaB2 construct, the PiggyBac transposase system was used as described previously (Odell *et al*., 2025). Antibiotic selection was performed using Puromycin at 3 µg/ml for at least 1 week, or until cells in a non-transformed well had all died. For stable expression of LBR-GFP, pseudovirus particles were made using 293TN cells as previously described (Earle *et al*., 2020). TKO MEFs were infected with virus twice on consecutive days, and on the third day hygromycin was added at 300 µg/ml for 1 week until all cells in a negative control well had died.

### Immunofluorescence

Cells (1 × 10^4^) were seeded on fibronectin-coated 12 mm glass coverslips overnight, then doxycycline (100 ng/ml) was added for 24 hours to induce expression of the exogenous lamin. Cells were fixed in 4% paraformaldehyde in PBS for 15 min at room temperature, followed by three 5-min washes with IF wash buffer containing 0.2% Triton X-100, 0.25% Tween 20 and 0.3% BSA in PBS. Cells were blocked in 3% BSA in PBS for 1 h, then primary antibodies were added for 1 h in blocking buffer at room temperature or overnight at 4°C. Primary antibodies used were: anti-FLAG (Sigma F7425, 1:1000), mouse anti-LBR (Abcam AB232731, 1:1000), mouse anti-lamin A/C (Santa Cruz sc-376248, 1:250), rabbit anti-lamin B1 (Abcam AB16048, 1:1000), and mouse anti-lamin B2 (Invitrogen 33-2100, 1:500).

DAPI was added 1:500 in PBS for 15 minutes to stain DNA. Secondary antibodies used were Alexa Fluor 488, 568, or 647-conjugated donkey anti mouse or rabbit-IgG antibodies (Invitrogen) diluted 1:250 in 3% BSA in PBS. Coverslips were mounted on glass slides using Mowiol and kept in the dark until imaging. Confocal immunostaining images were acquired on a Zeiss LSM900 series confocal microscope with Airyscan module using a 40× water immersion objective (NA 1.2). The optimal z-slice size was automatically determined using Zen Blue (Zeiss) software. Airy units for images were set between 1.5 and 2.5. Images are displayed as maximum intensity projections.

### Immunoblotting

For standard immunoblotting (Figure 1D and Figure 5A), 1 × 10^5^ cells were seeded in wells of a 6-well plate and allowed to adhere overnight, then doxycycline was added to induce expression of the exogenous lamin for 24 h. Cells were lysed in high-salt RIPA buffer containing 12 mM sodium deoxycholate, 50 mM Tris-HCl pH 8.0, 750 mM NaCl, 1% (v/v) NP-40 Alternative, 0.1% (v/v) SDS. Lysates were then vortexed for 5 min, sonicated (Branson 450 Digital Sonifier) for 30 s at 36% amplitude, and centrifuged at 4°C for 10 min at 14,000 g. Equal protein amounts were mixed with 4× Laemmli buffer (Biorad 1610747), denatured by boiling for 3 min, loaded onto 4–12% Bis-Tris gels (Invitrogen NP0322), run for 1.5 h at 100 V, then transferred for 1 h at 16 V onto PVDF membrane. Membranes were blocked for 1 h in blocking buffer containing 3% BSA in Tris-buffered saline plus 1% Tween 20 (TBST). Primary antibodies used: anti-FLAG (Sigma F7425, 1:2000) and anti LBR (Abcam AB232731, 1:1000). Primary antibodies were incubated with membranes overnight, then washed 3 times with TBST before addition of secondary antibodies.

For Phos-tag blotting, cells were seeded as above, except for roscovitine studies where 20 µM roscovitine or vehicle was added after 24 h of doxycycline addition and cells were incubated for an additional 24h prior to lysis. Cells were lysed on ice using 2× Laemmli buffer (Biorad 1610737) supplemented with 1 mM PMSF, 5% (v/v) β-mercaptoethanol, and phosphatase inhibitor tablet (Thermo Scientific A32957). Lysates were then vortexed for 5 min, sonicated (Branson 450 Digital Sonifier) for 30 s at 36% amplitude, and centrifuged at 4°C for 10 min at 14,000 g. Lysates were loaded onto precast SuperSep Phos-tag (50 µmol/L) gels (Fujifilm 192-18001) and run for 3 h at 100 V in a cold room maintaining a temperature of 4°C. Gels were washed 3 × 5 min in transfer buffer containing 0.025 mol/L Tris, 0.192 mol/L glycine, and 10 mmol/L EDTA pH 8.0. Gels were washed once more with transfer buffer without EDTA, and then proteins were transferred to PVDF membrane for 1.5 h at 16 V. Membranes were blocked and probed with anti-LBR antibodies as described above. Secondary antibodies used: Licor IRDye 680RD donkey anti-mouse-IgG (926-68072; 1:5000) and Licor IRDye 800CW Donkey anti-Rabbit IgG (926-32213; 1:5000). Secondary antibodies were added for 1 h at room temperature in blocking buffer, followed by three 10-min washes with TBST. Membranes were imaged using the Odyssey Licor scanner and then cropped and brightness and contrast was adjusted using Image Studio Lite (version 5.2) software. Band intensities were determined using the automatic band detection function of Image Studie Lite (version 5.2).

### Time-lapse microscopy

To observe localization of LBR-GFP over time, 1 × 10^5^ cells were seeded on fibronectin coated glass bottom 35-mm dishes overnight prior to experiment.

Timelapse imaging was performed using a Keyence BZ-X810 microscope with 40 × Plan Apochromat lens (NA 0.95) and GFP filter cube. Cells were maintained in a humidified environment with temperature control set to 37°C and 5% CO_2_. Image acquisition was automated at 20-minute intervals.

### Fluorescence recovery after photobleaching (FRAP)

For FRAP experiments, 1 × 10^5^ cells were seeded on a fibronectin-coated 35 mm glass-bottom dish the day before transfection. Transient transfection of full-length LBR-GFP was achieved using Mirus TransIT-X2 transfection reagent according to manufacturer’s instructions. 100 ng/ml doxycycline was added simultaneously with transfection reagent to induce expression of the desired lamin construct. Cells were allowed to express LBR-GFP and the exogenous lamin for 24 hours prior to FRAP experiments. Photobleaching was performed using Zeiss LSM 900 confocal microscope with a 40 × water immersion objective (NA 1.20) using the “iterative bleaching” function. The focus was set to the central z-position for each cell, such that the LBR-GFP nuclear rim was visible. A ROI at the nuclear envelope was drawn manually for each cell. Three frames were acquired before photobleaching, then LBR-GFP was bleached using 100% 488 laser power at maximum scan speed (corresponding to a pixel dwell time of 0.687 µs) for 10 iterations within the ROI, followed by image acquisition every 0.5 s for 200 s following photobleaching. Efforts were made to select cells for photobleaching with comparable LBR-GFP expression levels and similarly sized nuclear envelope ROIs for bleaching were drawn across conditions. FRAP analysis was performed as described (Odell *et al*., 2026). Briefly, mean fluorescence intensity of LBR-GFP within the bleached regions was determined for all frames of the timelapse. The mean fluorescence intensity of a background region devoid of cells was subtracted from the fluorescence intensity measurements, and data were normalized to the total cell fluorescence intensity at each time point to control for general photobleaching as a result of continuous imaging. Data were then normalized to the average pre-bleach intensity to yield the normalized intensity over time within the bleached region. FRAP data from 20-30 cells were averaged to yield the FRAP recovery curves, and data were fit to a single variable exponential curve using Microsoft Excel.

### Image analysis

For measurements of nuclear-to-cytoplasmic ratios of LBR, nuclear LBR levels were measured using a previously described FIJI macro (Odell *et al*., 2024). Briefly, immunofluorescence images were thresholded based on the DAPI staining to determine the nuclear region of interest (ROI), and the mean intensity in the LBR channel within the nucleus was measured. Then, mean cytoplasmic LBR levels were obtained by manually drawing a ROI adjacent to the corresponding nucleus for each cell. Mean nuclear LBR intensity was divided by mean cytoplasmic LBR intensity for each cell. Intensity profile measurements were performed using a FIJI macro available on request. Briefly, this macro used the ‘Plot Profile’ feature in FIJI software to measure the LaB1 or LaB2 intensity across a line drawn across a z-slice through the center of the nucleus. To account for differences in nuclear size, the intensity profiles are converted into relative nuclear distances by measuring the average intensity in each of 50 equally sized bins. For all analysis, cells on the edges of the image, dead cells, or mitotic cells were excluded manually from the analysis.

### Statistical analysis and figure generation

All analyses were performed using GraphPad Prism. Information on statistical tests used and significance values are present in each figure caption. Our statistical analysis was developed in close consultation with the Cornell Statistical Consulting Unit. Figures were assembled using Adobe Illustrator.

## Acknowledgements

We thank Loren Fong (UCLA) for generously sharing the mLmnb2-pCMV plasmid and Ohad Medalia (University of Zurich) for sharing the DARPin expression constructs. We thank the Biotechnology Resource Center (BRC) Flow Cytometry Facility (RRID: SCR_021740) and sequencing facility (RRID: SCR_021727) at the Cornell Institute of Biotechnology for their resources and technical assistance. This work was supported by awards from the Volkswagen Foundation (A130142 to J.L.), the National Institutes of Health (R01 HL082792, R01 GM137605, R35 GM153257, and R01 AR084664 to J.L.), the National Science Foundation (URoL2022048 to J.L.), and the Leducq Foundation (20CVD01 and 24CVD03 to J.L.). The content of this manuscript is solely the responsibility of the authors and does not necessarily represent the official views of the National Institutes of Health.

## Data availability stahtement

The authors confirm that the data supporting the findings of this study are available within the article and its supplementary materials.

