## Supplementary Materials for "Antagonistic contributions of A-type and B-type lamins to LBR localization and dynamics"

**Supplemental Figures:**

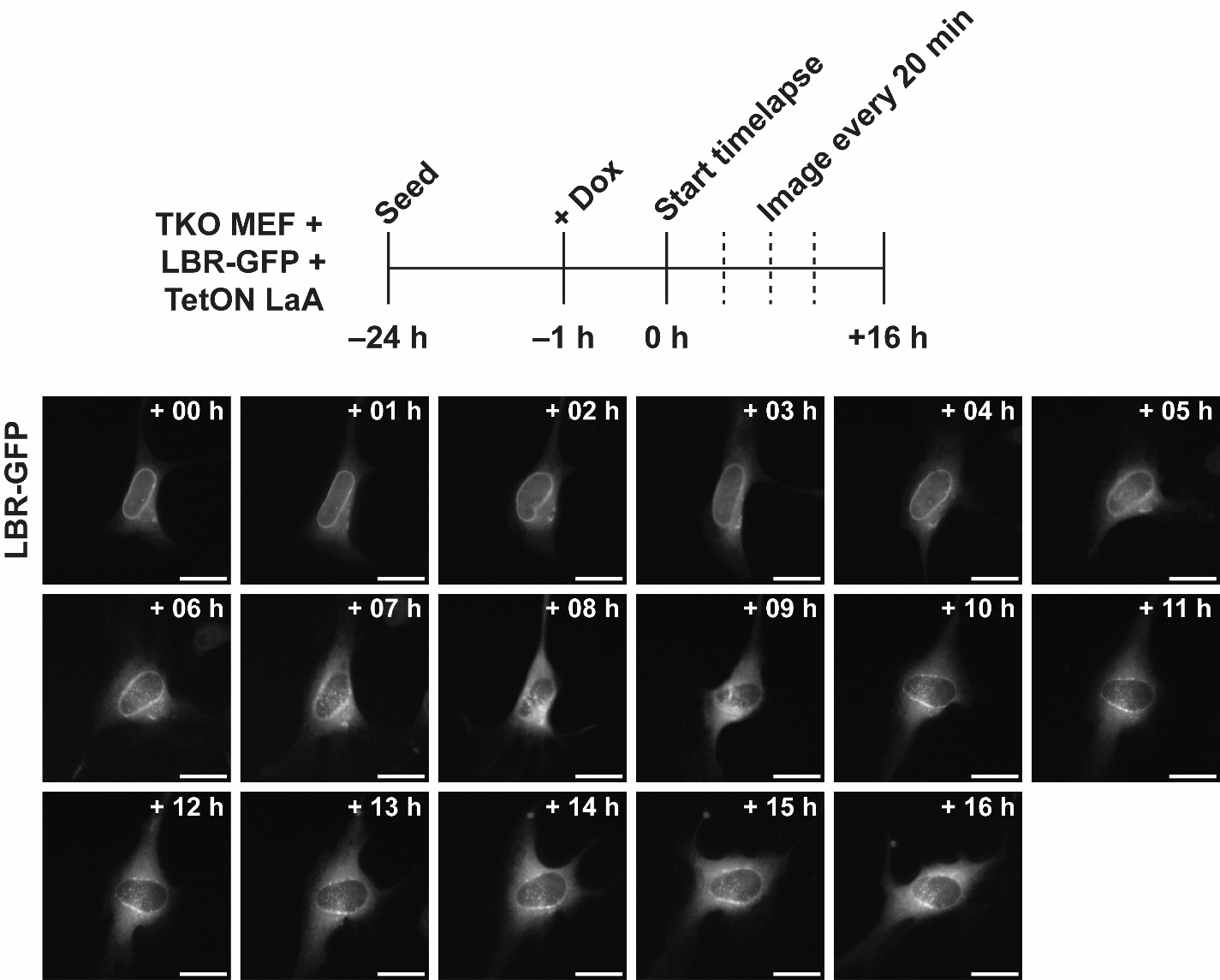

**Supplemental Figure 1: LaA induces LBR-GFP displacement in TKO MEFs during interphase.** (**A**) Overview of experimental setup. Cells were seeded on glass-bottom dishes 24 hours prior to the start of the time-lapse experiment and allowed to adhere overnight. An hour prior to the beginning of image acquisition, doxycycline was added to cells to induce LaA expression, and cells were transferred to the microscope chamber to allow for temperature equilibration. Then, frames were taken every 20 minutes to acquire dynamic behaviors of LBR-GFP over time. (**B**) Representative frames for an image series described in panel A. Scale bar = 20 µm.

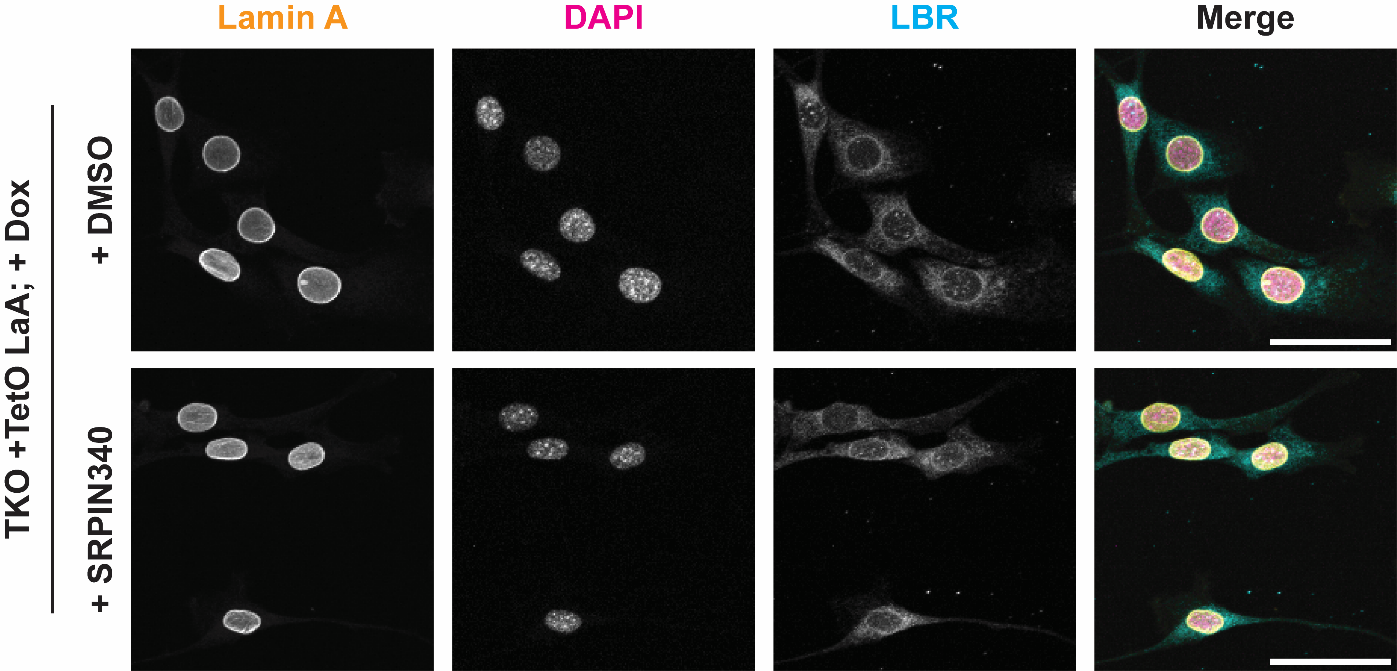

**Supplemental Figure 2: SRPIN340 does not prevent LaA-induced LBR displacement in TKO MEFs.** Cells were first induced to express LaA by adding doxycycline to the media for 24 hours. SRPIN340 or vehicle (DMSO) was added to cells at a final concentration of 10 µM for an additional 24 hours prior to fixation and immunofluorescence staining. Scale bar = 50 µm.

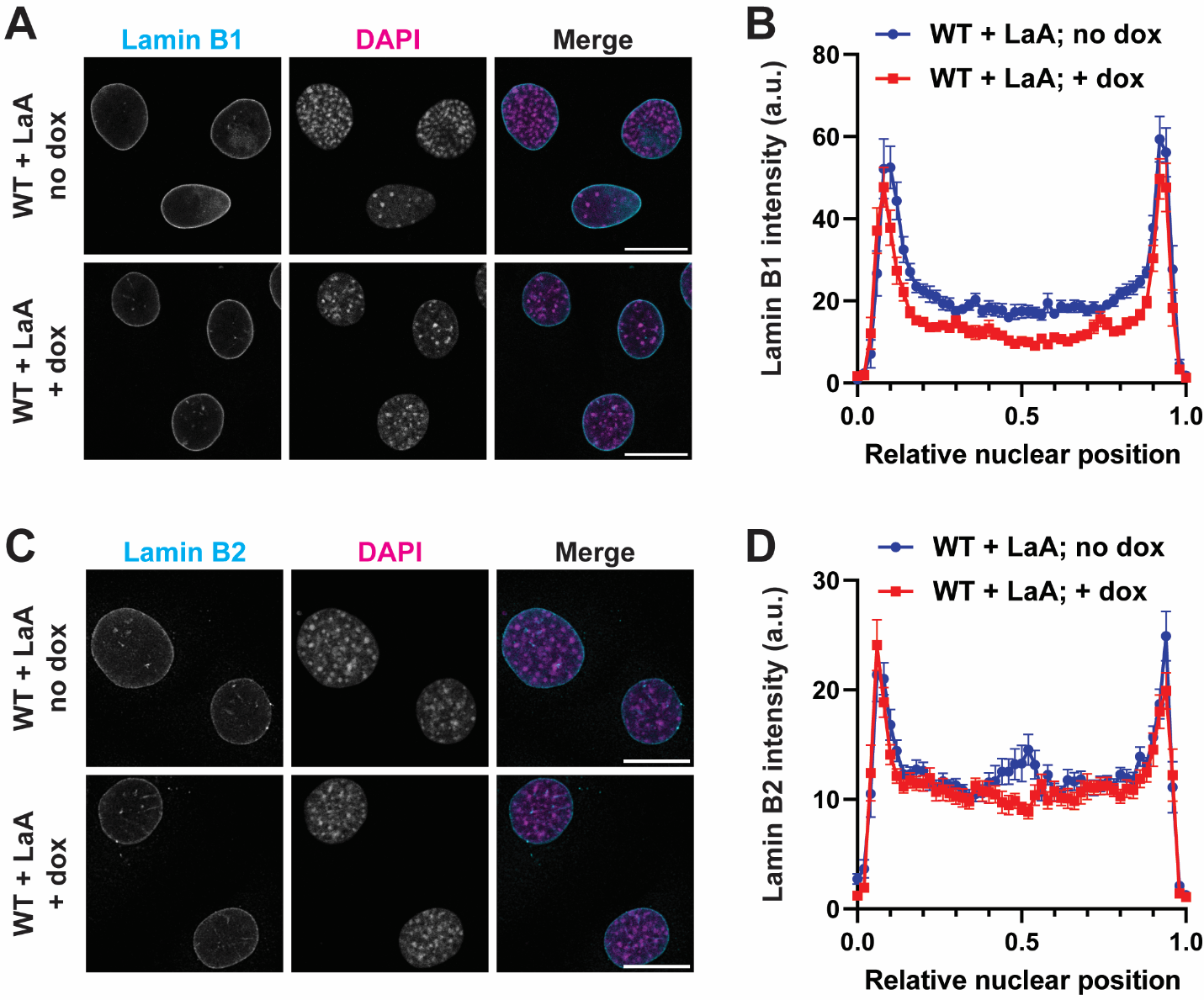

**Supplemental Figure 3: Localization of B-type lamins in wild-type MEFs overexpressing LaA. (A**) Immunofluorescence labelling for endogenous LaB1 in wild-type MEFs expressing wild-type (WT) LaA. Scale bar = 10 µm. (**B**) Intensity profiles of LaB1 measured by drawing a line through the midplane of the nucleus. Peaks at the edges of the plot represent the nuclear envelope signal. Points and error bars indicate mean ± s.e.m. (**C**) Immunofluorescence labelling for endogenous LaB2 in wild-type MEFs expressing WT LaA. Scale bar = 10 µm. (**D**) Intensity profiles of LaB2 measured by drawing a line through the midplane of the nucleus. Peaks at the edges of the plot represent the nuclear envelope signal. Points and error bars indicate mean ± s.e.m.

**Supplemental Table**

| Primer Name | Sequence 5’-3 | Use |
| --- | --- | --- |
| LaminB2_fwd | accctcgtaaaggtctagagaccatggactacaaagacgatgacgacaagatgagcgcgccgcattcg | Cloning FLAG-LaB2 |
| LaminB2_rev | ccgtttaaactcattactaatcacatcagtcggcagcc | Cloning FLAG-LaB2 |
| LBR-GFP_fwd | ctcgagactagttctagagcatgccaagtaggaaatttg | Cloning LBR-GFP |
| LBR-GFP_rev | ggggggagggagaggggcggttacttgtacagctcgtc | Cloning LBR-GFP |

**Supplementary Table 1:** List of primers used for Gibson cloning the indicated constructs.
